# Synergistic and independent roles for Nodal and FGF in zebrafish CPC migration and asymmetric heart morphogenesis

**DOI:** 10.1101/2024.01.05.574380

**Authors:** Vanessa Gonzalez, Meagan G. Grant, Makoto Suzuki, Briana Christophers, Jessica Rowland Williams, Rebecca D. Burdine

## Abstract

Asymmetric development of the vertebrate heart is driven by a complex sequence of morphogenetic cell movements, coordinated through precise interpretation of signaling cues by the heart primordia. Here, we show that Nodal signaling functions synergistically with FGF to stimulate the migration of cardiac progenitor cells (CPCs) during cardiac jogging—the first morphological asymmetry observed in zebrafish heart development. While Nodal directs the asymmetric migration of CPCs, we find FGF signaling to be dispensable for this asymmetry, suggesting that FGF plays a permissive rather than instructive role. We further demonstrate that Nodal signaling induces asymmetries in actin cytoskeletal dynamics that correlate with the directional migration of CPCs, while FGF does not influence this actin asymmetry. In addition to influencing jogging, FGF and Nodal synergize to ensure proper heart looping. We also provide evidence that FGF contributes to heart looping by promoting the differentiation of the second heart field. Together, these findings offer insight into how the spatiotemporal dynamics of signaling pathways regulate the cellular behaviors driving organ morphogenesis.

**Summary statement:** This study explores the synergistic and independent roles of Nodal and FGF signaling in generating heart asymmetry.

## Introduction

Vertebrate heart development requires the faithful execution of intricate and successive cell movements. The coupling of these movements to morphogenesis is mediated by the precise interpretation of signaling cues by the heart primordia. Perturbations in these signaling pathways can underlie cellular processes that go awry in the roughly 40,000 infants born with a congenital heart defect (CHD) each year in the United States (Tsao et al., 2022). Therefore, a comprehensive analysis of signaling cues that direct cardiac cell behavior is critical for understanding the pathogenic mechanisms of CHDs.

The presence of conserved molecular mechanisms, the amenability to genetic manipulation and live imaging, and the ability for continued embryonic development in the presence of heart defects make zebrafish particularly advantageous for studying the cellular behaviors underlying cardiac morphogenesis (Bakkers, 2011; Genge et al., 2016; Grant et al., 2017; Nguyen et al., 2008; Smith and Uribe, 2021). During the initial stages of zebrafish heart development, cardiac progenitor cells (CPCs) are organized bilaterally within the anterior lateral plate mesoderm (LPM) as epithelial sheets, with ventricular progenitor positioned medial to atrial cells. These sheets converge on the embryonic midline, where they form the cardiac cone, a disc-shaped structure comprised of atrial cells at its base and ventricular cells at its apex (19 hours post fertilization, hpf; Bakkers, 2011; Grant et al., 2017; Smith and Uribe, 2021).

During a process known as cardiac jogging, CPCs migrate in a leftward and anterior direction, with cells on the left half of the cardiac cone migrating more rapidly than the right-sided cells (20-24 hpf). This asymmetry in migration velocities results in clockwise rotation of the cardiac cone and displacement of atrial cells to the left and anterior of ventricular cells as the heart tube elongates. Cardiac jogging culminates in the asymmetric placement of the heart tube to the left of the embryonic midline, establishing the first asymmetry in the heart (Baker et al., 2008; de Campos-Baptista et al., 2008; Kidokoro et al., 2022; Lenhart et al., 2013; Rohr et al., 2008; Smith et al., 2008; Veerkamp et al., 2013). Post jogging, the second heart field (SHF) is progressively added to the poles of the heart tube and contributes to heart tube elongation, eventually forming the outflow tract (24-48 hpf) (de Pater et al., 2009; Hami et al., 2011; Zhou et al., 2011). Between 30-48 hpf, the heart undergoes an evolutionarily conserved process known as cardiac looping. During this process, the heart tube bends rightward to begin defining and aligning the atrial and ventricular chambers of the heart, with actin polymerization being critical for dictating looping chirality (Desgrange et al., 2018; Noel et al., 2013; Smith and Uribe, 2021). Along with cardiac looping, the cardiac chambers undergo expansion and become morphologically discernible from one another via cardiac ballooning. Regionalized differences in cell morphology bring about chamber formation, and blood flow and contractility are critical for regulating these morphologies (Auman et al., 2007; Dietrich et al., 2014; Smith and Uribe, 2021).

Left-sided Nodal signaling in the LPM governs asymmetric behaviors during jogging and dextral looping of the heart. Nodal directs the asymmetric migration of CPCs during cardiac jogging by increasing the velocities of left-sided cells in the cardiac cone and properly positioning the heart to the left of the embryonic midline (Baker et al., 2008; de Campos-Baptista et al., 2008; Kidokoro et al., 2022; Lenhart et al., 2013; Rohr et al., 2008; Veerkamp et al., 2013). While looping of the jogged heart tube involves intrinsic mechanisms (Noel et al., 2013), this process is influenced by earlier Nodal signaling to produce robustness in dextral looping morphogenesis (Grimes et al., 2020).

Although Nodal signaling is the dominant signal for sidedness, it is well-established that Nodal signals integrate with other signaling pathways to orchestrate events controlling organogenesis (Hill, 2016; Massague, 2003). For example, during cardiac cone rotation and subsequent jogging, Nodal and BMP signals act synergistically to influence cell migration (Lenhart et al., 2013; Veerkamp et al., 2013). How other signaling pathways may integrate with Nodal signaling to orchestrate asymmetric heart morphogenesis remains unknown.

Here, we characterize the synergistic and independent functions of Nodal and FGF signaling during asymmetric morphogenesis of the zebrafish heart. We find FGF and Nodal both stimulate CPC migration during cardiac jogging, but only Nodal is involved in directing asymmetric cell migration suggesting that FGF plays a permissive role in this process. Consistent with these results, only Nodal induces asymmetries in actin cytoskeletal dynamics that correlate with directional migration of the CPCs; FGF signaling seems to have no influence on actin asymmetry in the CPCs. Later in development, FGF and Nodal signals synergize to promote proper heart looping. We provide evidence that the role of FGF in cardiac looping may be through promoting SHF addition to the ventricle, suggesting the SHF promotes proper looping *in vivo*. Together, these findings shed insight into how Nodal and FGF signals function to ensure proper migration and morphogenesis in the heart at multiple stages.

## Results

### FGF and Nodal signals function synergistically during heart tube formation and jogging

Previous studies have shown that the left-sided activation of Nodal in the anterior LPM drives the directional migration of CPCs, determining the direction of heart tube displacement from the embryonic midline during cardiac jogging (Baker et al., 2008; de Campos-Baptista et al., 2008; Kidokoro et al., 2022; Lenhart et al., 2013; Rohr et al., 2008; Veerkamp et al., 2013). To gain insight into how Nodal influences left-right asymmetries in CPC migration, we utilized our previous microarray studies that identified genes upregulated in Nodal-positive CPCs that were downregulated in Nodal-negative CPCs (Williams, 2015). This analysis identified several components of the FGF signaling pathway, including FGF receptors *fgfr2*, *fgfr3*, and *fgfr4* and transcriptional targets of the pathway *erm*, *pea3*, and *spry4*. FGF signaling components have been identified as Nodal targets during gastrulation in zebrafish, including *fgf17b*, *fgf3*, and *fgf8* (Bennett et al., 2007). In zebrafish, *fgf8* is required for the development of CPCs and is expressed in the cardiac cone prior to cardiac jogging (Fig. 1A; Reifers et al., 1998; Reifers et al., 2000). Given the recognized importance of FGF signaling in heart development and the potential connection to Nodal, we tested whether FGF signaling contributed to heart jogging and looping, the two major left-right asymmetric events of the heart.

**Figure 1.**
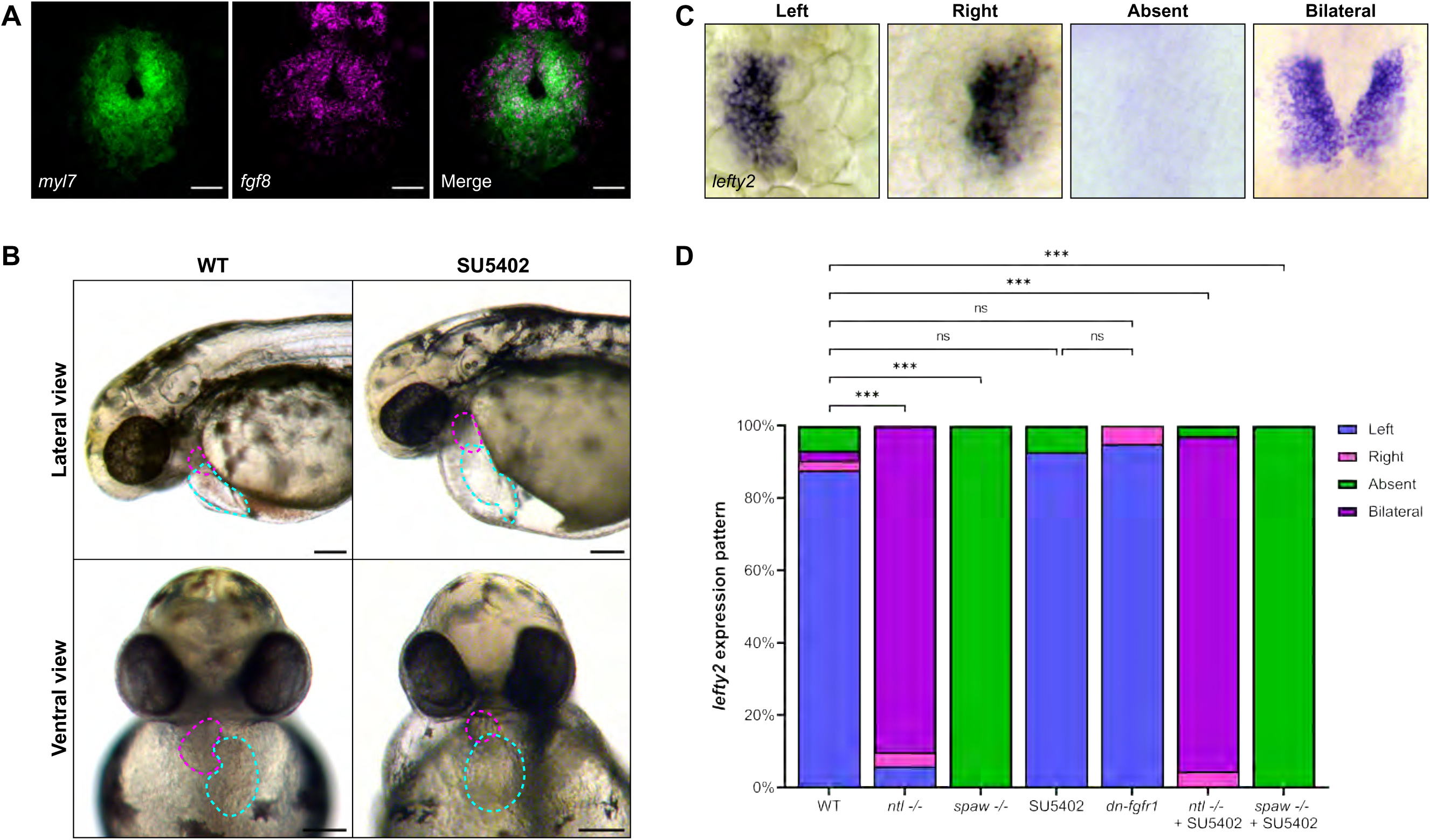
FGF signaling is active in the developing heart and necessary for proper heart development. (**A**) RNA *in situ* hybridization (ISH) by hybridization chain reaction (HCR) for *fgf8* (magenta) and *myl7* (green) in the cardiac cone of a 20 hours post-fertilization (hpf) wild-type (WT) embryo. Scale = 25 µm. (**B**) Representative images of 48 hpf WT embryos or embryos treated with FGF inhibitor SU5402 from 19-30 hpf (ventricle in magenta, atrium in cyan). SU5402-treated embryos display aberrant cardiac looping, misshapen and misplaced chambers, and pericardial edema. (**C**) ISH for Nodal target gene *lefty2* (*lft2*) in the cardiac cone of 20 hpf embryos shows correct left-sided *lefty2* and incorrect right, bilateral, and absent *lefty2* expression. (**D**) Quantification of *lefty2* expression sidedness. n is WT = 74, *ntl^-/-^*= 50, *spaw^-/-^* = 36, SU5402 = 41, *dn-fgfr1* = 20, *ntl^-/-^*+ SU5402 = 177, *spaw^-/-^* + SU5402 = 63. Statistical significance was determined by Chi-square analysis. (**A, C**) Images are dorsal views, with anterior to the top and left to the reader’s left. ns = not significant, *** = p<0.0001.

Since early inhibition of FGF signaling affects cardiac progenitor development (Felker et al., 2018; Marques et al., 2008; Reifers et al., 2000), we cannot make use of mutants such as *fgf8/ace* or global knockdown techniques to study later cardiac phenotypes. Thus, to inhibit FGF signaling with precise temporal control, we administered the pan-FGFR inhibitor SU5402 (Mohammadi et al., 1997). The addition of SU5402 from 19-30 hpf resulted in various cardiac defects, including severe pericardial edema and aberrant chamber placement, phenotypes associated with errors in the asymmetric development of the heart, suggesting FGF signaling could be involved in this process (Fig. 1B).

To confirm that inhibiting FGF signaling with SU5402 did not affect the establishment and expression of left-right patterning genes, we analyzed expression of the Nodal target gene *lefty2* in the cardiac cone. *lefty2* expression serves as both a readout of Nodal responsiveness and a direct readout of the sidedness of Nodal in the embryo (Fig. 1C). In WT embryos, *lefty2* expression is restricted to the left side of the cardiac cone (Fig. 1D). *southpaw* (*spaw*) mutants, which lack Nodal expression in the LPM, lack *lefty2* expression (Grimes et al., 2020; Noel et al., 2013), while the majority of *ntl* (*ntl/ntla/tbxta*) mutants, which have bilateral Nodal expression in the LPM, express *lefty2* bilaterally (Amack and Yost, 2004), as expected (Fig. 1D). Loss of FGF signaling in SU5402-treated WT embryos did not alter the left-sidedness of Nodal target *lefty2*, nor did loss of FGF change the expression profile of *lefty2* in *spaw* and *ntl* mutants (Fig. 1D). To corroborate our results with SU5402, we used the *Tg(hsp70l:dnfgfr1-EGFP)* heat shock inducible transgenic line to inhibit FGF signaling and found this similarly did not alter *lefty2* expression (Fig. 1D). These results suggest that any effect of FGF on asymmetric heart development is downstream of left-right axis establishment.

To investigate the role of FGF signaling during cardiac jogging—the first morphological sign of asymmetry in the embryo—we inhibited FGF using SU5402 or heat shock–induced dnFGFR1 from 19 hpf at cardiac cone formation, and continued treatment through jogging and heart tube formation (Fig. 2A). Embryos were fixed at 26.5 hpf and the laterality of heart jogging was analyzed (Fig. 2B). In WT embryos, heart tubes are displaced to the left of the embryonic midline, producing the expected “left jog” (Fig. 2C). Both *spaw* mutants and *ntl* mutants exhibit primarily “midline jog”, where the heart tube fails to be displaced to either side of the midline (Fig. 2C). Inhibiting FGF signaling in WT, *spaw* mutant, and *ntl* mutant embryos with either the pharmacological or heat shock approach did not affect laterality, as hearts jogged similarly to respective WT and mutant controls (Fig. 2C).

**Figure 2.**
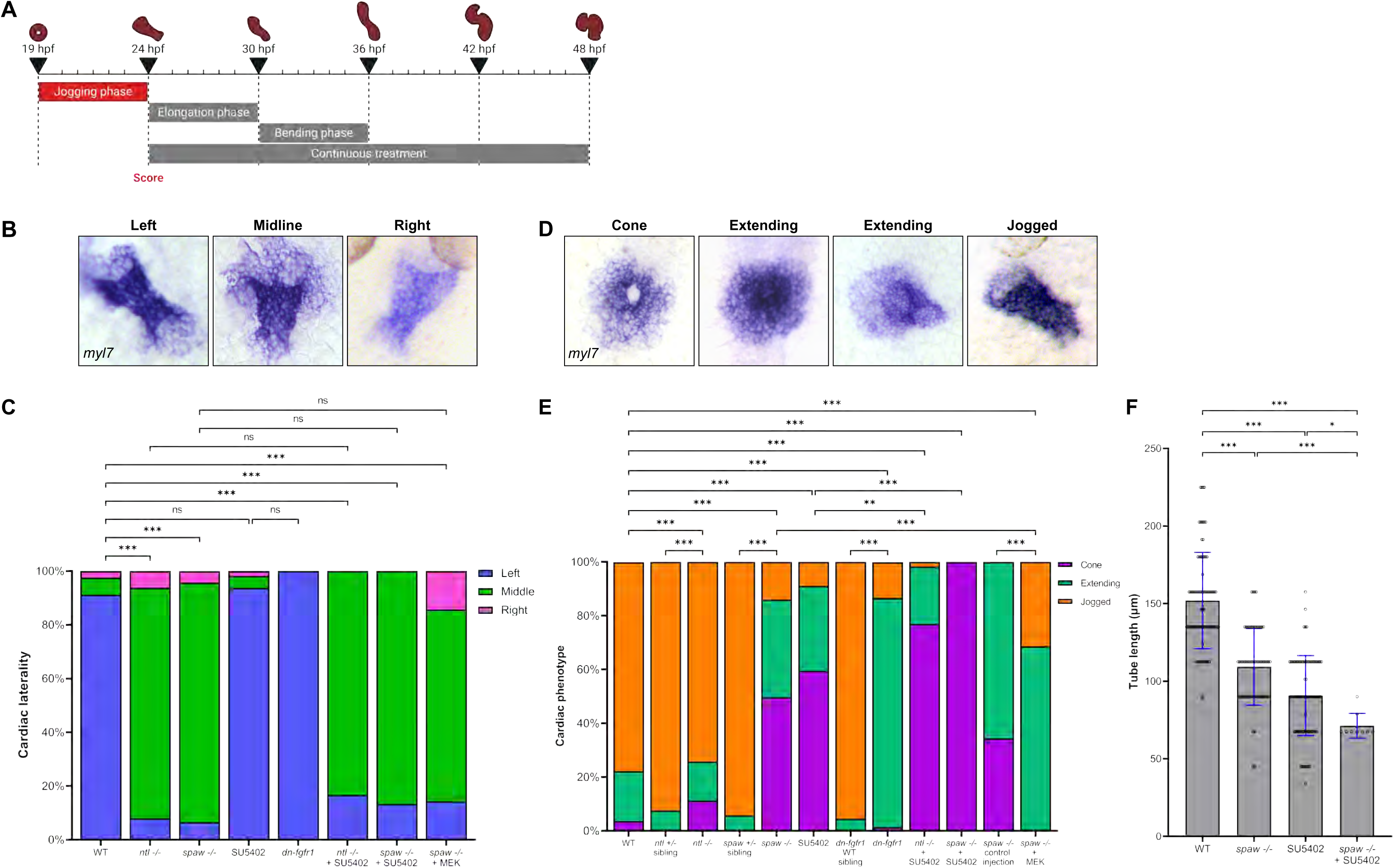
FGF signaling is necessary for proper CPC migration but not cardiac laterality during jogging. (**A**) Schematization depicting the 19-24 hpf SU5402 treatment window during the “Jogging phase”. (**B**) RNA ISH for *myl7* at 26.5 hpf visualizing correct left-sided positioning and incorrect midline and right-sided positioning of the heart tube. (**C**) Quantification of cardiac laterality phenotypes. n is WT = 534, *ntl^-/-^* = 113, *spaw^-/-^* = 138, SU5402 = 112, *dn-fgfr1* = 75, *ntl^-/-^* + SU5402 = 12, *spaw^-/-^* + SU5402 = 30, *spaw^-/-^* + MEK = 14; Chi-square analysis. (**D**) RNA ISH for *myl7* at 24 hpf displaying the stages from cone through extending to correctly jogged heart tubes we see in our treated embryos. (**E**) Quantification of cardiac migration phenotypes. Number of embryos (n) is WT = 686, *ntl^+/-^* sibling = 213, *ntl^-/-^*= 152, *spaw^+/-^* sibling = 87, *spaw^-/-^* = 171, SU5402 = 531, *dn-fgfr1* WT sibling = 66, *dn-fgfr1* = 225, *ntl^-/-^*+ SU5402 = 117, *spaw^-/-^* + SU5402 = 146, *spaw^-/-^* control injection = 105; *spaw^-/-^* + MEK = 35; Chi-square analysis. (**F**) Quantification of heart tube length at 26.5 hpf. n is WT = 84, *spaw^-/-^* = 62, SU5402 = 80, *spaw^-/-^* + SU5402 = 9. Statistical significance determined by Student’s t-test. (**B, D**) Images are dorsal views, with anterior to the top and left to the reader’s left. ns = not significant, * = p<0.05, ** = p<.001, *** = p<0.0001.

While the previous results suggest that FGF signaling does not influence jogging laterality, we observed that the inhibition of FGF signaling led to fewer scorable hearts due to delayed tube formation, indicating a potential effect on CPC cell migration. To determine if FGF signaling affects cell migration during jogging, we inhibited FGF signaling at 19 hpf and assessed embryos at 24 hpf, a time when most wild-type (WT) embryos have completed jogging (Fig. 2D). *ntl* mutants exhibit only a slight delay in cone-to-tube transition, compared to WT (Fig. 2E). By contrast, *spaw* mutants display a significant delay in tube formation, with most hearts remaining at cone stage or starting to extend at 24 hpf (Fig. 2E). Both SU5402-treated and heat-shocked dnFGFR1 embryos display significant delays in tube formation (Fig. 2E). Interestingly, treating *ntl* and *spaw* mutants with SU5402 further increased delays in tube formation, with most hearts remaining in the cardiac cone stage (Fig. 2E). The strongest effect was observed in *spaw* mutants treated with SU5402, in which all embryos analyzed remained at the cardiac cone stage at 24 hpf (Fig. 2E). Given this is a timing-based phenotype, we compared *ntl* mutants, *spaw* mutants, and heat-shocked dnFGFR1 embryos to their heterozygous/WT siblings and found each pair to be significantly different from each other (Fig. 2E). Taken together, these results suggest that both the Nodal and FGF pathways promote CPC migration during jogging, leading to the formation of the heart tube at the appropriate time.

To determine whether activating signaling downstream of FGF could restore CPC migration in *spaw* mutants, we injected these mutants with photoswitchable MEK (psMEK). psMEK is active when exposed to 500 nM light, activating the MAPK-ERK pathway, which functions downstream of receptor tyrosine kinases, including FGFR (Patel et al., 2019). When psMEK is activated during jogging, a significantly higher number of embryos develop jogged and extended heart tubes compared to untreated *spaw* mutants (Fig. 2E). However, jogging laterality was not rescued (Fig. 2C), suggesting that while MAPK-ERK signaling enhances overall cell motility, it does not influence laterality.

While heart tube formation was delayed at 24 hpf in FGF-deficient and Nodal-deficient embryos, heart tubes eventually formed in all groups after several additional hours but appeared shorter than those in WT. To quantify this observation, we inhibited FGF during jogging and then measured heart tube length at 26.5 hpf in embryos where the heart tube had completed jogging. Heart tubes were significantly shorter in SU5402-treated embryos, possibly due to delays in tube elongation or effects on the SHF as reported previously (Fig. 2F; de Pater et al., 2009; Felker et al., 2018; Lazic and Scott, 2011). Heart tubes were also shorter in *spaw* mutants, suggesting that tube elongation is delayed or that elongating in a midline position may be constrained by surrounding tissues (Fig. 2F). Combined loss of FGF and Nodal signaling (SU5402-treated *spaw* mutants) was additive, producing significantly shorter heart tubes than *spaw* mutants or SU5402-treated embryos (Fig. 2F). Overall, these data suggest that Nodal and FGF signaling synergize to promote cell migration during heart tube formation and jogging, while only Nodal signaling is required to induce lateralized migration during this process.

### Nodal and FGF signals regulate the velocity dynamics and directionality of CPCs during heart tube formation

Perturbation of the Nodal and FGF pathways results in defects in heart tube morphogenesis that we hypothesize are due to defects in cell migration. To explore this possibility further, we conducted confocal live imaging of embryos that express EGFP under the cardiac-specific *myl7* promoter (Huang et al., 2003) to quantify metrics of cell migration during jogging (Fig. 3A). Starting from the formation of the cardiac cone, cell movements were analyzed for three hours in each condition. WT embryos completed jogging within the three-hour time frame and formed a left-lateralized heart tube (Fig. 3B, Movie 1). *ntl* mutants completed jogging in the same time frame but produced midline heart tubes due to bilateral Nodal expression (Fig. 3B, Movie 2). *spaw* mutants also displayed midline heart tubes; however, these CPCs migrated more slowly compared to WT or *ntl* mutants, often resulting in incomplete tube extension during the three-hour time frame (Fig. 3B, Movie 3). Similarly, CPCs in SU5402-treated WT embryos migrated more slowly compared to WT or *ntl* mutants, resulting in incomplete tube extension during the imaging time frame (Fig. 3B, Movie 4). The effects of Nodal and FGF signaling on migration are additive, as CPCs in SU5402-treated *spaw* mutants scarcely migrated during the same time (Fig. 3B, Movie 5).

**Figure 3.**
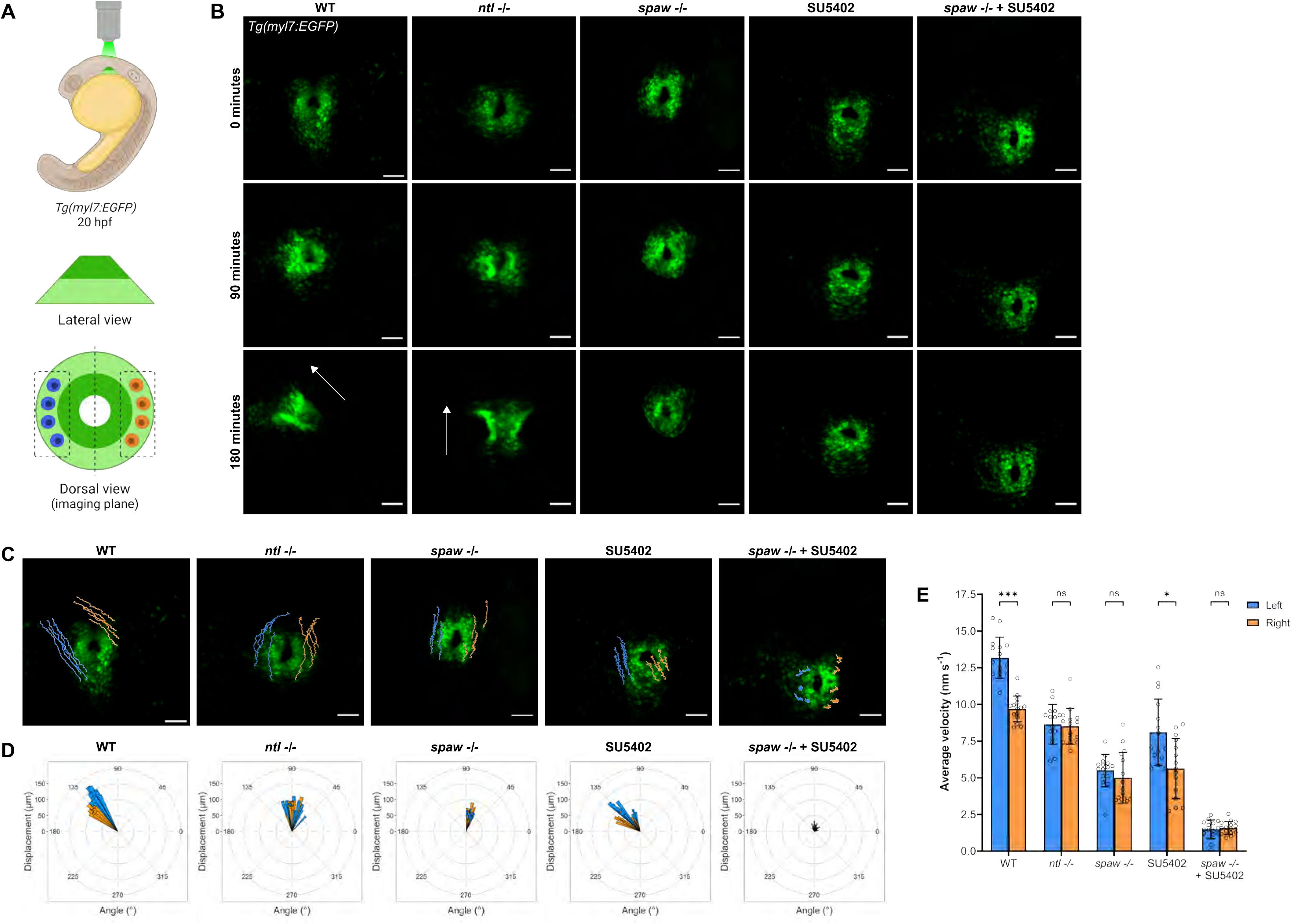
FGF signaling promotes CPC migration while Nodal promotes CPC migration and laterality during jogging. (**A**) Schematization of imaging strategy for time-lapse imaging of the cardiac cone throughout jogging, with outer atrial cells in light green and inner ventricular cells in dark green. Left-sided (blue) and right-sided (orange) cells that underwent tracking for analysis are shown. (**B**) Representative time-lapse images depicting cardiac progenitor cell (CPC) migration from the formation of the cardiac cone through the next 3 hours of development. WT and *ntl^-/-^* embryos typically complete jogging and tube formation in this timeframe. Arrows indicate heart tube direction. Scale = 50 µm. (**C**) Trajectories of left-sided (blue) and right-sided (orange) CPCs during 3 hours of jogging. Scale = 50 µm. (**D**) Rose plots demonstrating the angle of displacement of each tracked left-sided (blue) and right-sided (orange) CPC after 3 hours of jogging. (**E**) Average velocity of left-sided and right-sided CPCs during 3 hours of jogging; Student’s t-test. (**C-E**) n = 3 embryos (5 left-sided and 5 right-sided cells tracked per embryo) per condition. (**B-C**) Images are dorsal views, with anterior to the top and left to the reader’s left. ns = not significant, * = p<.05, *** = p<0.0001.

To characterize asymmetric migration during jogging, we quantified the trajectories of the outermost atrial CPCs, which display the largest disparity in left-right migration rates (Lenhart et al., 2013) (Fig. 3C). Analysis of the direction and magnitude of CPC trajectories revealed that CPCs in WT embryos exhibit a marked anterior and leftward displacement, while CPCs in *ntl* and *spaw* mutants fail to undergo strong left or right displacement (Fig. 3D). Furthermore, in *ntl* and *spaw* mutants, overall anterior displacement of CPCs was decreased (Fig. 3D). SU5402-treated embryos display a leftward bias in CPC migration, though the overall displacement was not as marked as in WT embryos, and anterior migration was reduced in these embryos (Fig. 3D). Both lateral and anterior migrations are disrupted in SU5402-treated *spaw* mutants, further confirming that Nodal and FGF signals work synergistically to promote CPC migration during jogging (Fig. 3D). The CPC trajectories further suggest that Nodal and FGF signals contribute to the general anterior migration of the heart during jogging, while asymmetric Nodal signals direct lateral migration.

To better understand the contributions of Nodal and FGF signals to cell migration, we analyzed the migration velocities of CPCs. Cardiac laterality is known to be directed by left-right biases in cell migration velocity driven by Nodal signaling, with left CPCs displaying a higher velocity than right CPCs in WT embryos (de Campos-Baptista et al., 2008; Kidokoro et al., 2022; Lenhart et al., 2013; Veerkamp et al., 2013). Given the bilateral exposure to Nodal of CPCs in *ntl* mutants, we expected migration to be increased on both sides of the cardiac cone compared to WT, as we observed in *ntl* morphants (Lenhart K. F. et al., 2013). However, we found that *ntl* mutants did not display increased migration velocities compared to WT, although the *ntl* mutant velocities were higher than those in *spaw* mutants. As expected, the migration rates of left and right CPCs in *ntl* mutants were symmetrical (Fig. 3E).

Consistent with previous findings (de Campos-Baptista et al., 2008; Kidokoro et al., 2022; Lenhart et al., 2013; Veerkamp et al., 2013), migration velocities of CPCs in *spaw* mutants are symmetric and reduced compared to WT (Fig. 3E). Treatment with SU5402 in WT embryos produces a pronounced reduction of migration velocity similar to that of *spaw* mutants (Fig. 3E). However, asymmetry in left versus right migration velocities is still present, as expected given that heart tubes jog correctly to the left in these embryos (Fig. 1D, 2C, 2F, 3E). Intriguingly, SU5402-treated *spaw* mutants display a more severe reduction in migration velocity compared to *spaw* mutants or SU5402-treated embryos, further suggesting synergy between Nodal and FGF signals in this process (Fig. 3E). Taken together, our findings on CPC migration, velocity, and trajectories all support Nodal acting instructively to bias left migration velocities, while FGF acts permissively and synergistically with Nodal to promote overall CPC velocities during heart tube morphogenesis.

### Nodal induced asymmetries in actin polymerization correlate with left-right asymmetries in CPC migration

Cell migration is generally driven by dynamic assembly and disassembly of actin filaments (Schaks et al., 2019). Since Nodal signaling can regulate actin during cardiac looping and other developmental processes (Noel et al., 2013; Woo et al., 2012), we hypothesized that Nodal-driven asymmetric migration of CPCs during jogging may also result in asymmetric actin dynamics. To assess this possibility, we utilized a *Tg*(*myl7:Lifeact-EGFP*) transgenic line, in which EGFP-labeled filamentous actin (F-actin) is restricted to CPCs (Huang et al., 2003; Reischauer et al., 2014; Riedl et al., 2008), permitting us to observe F-actin levels throughout jogging (Fig. 4A). Although CPCs migrate while connected to their neighbors in an epithelial state (Trinh and Stainier, 2004), we observe dynamic protrusions from individual cells during jogging (Fig. 4B, Movie 6). To our knowledge, this is the first time the actin-rich protrusions of CPCs have been captured live during jogging, demonstrating the strong migratory capacity of CPCs.

**Figure 4.**
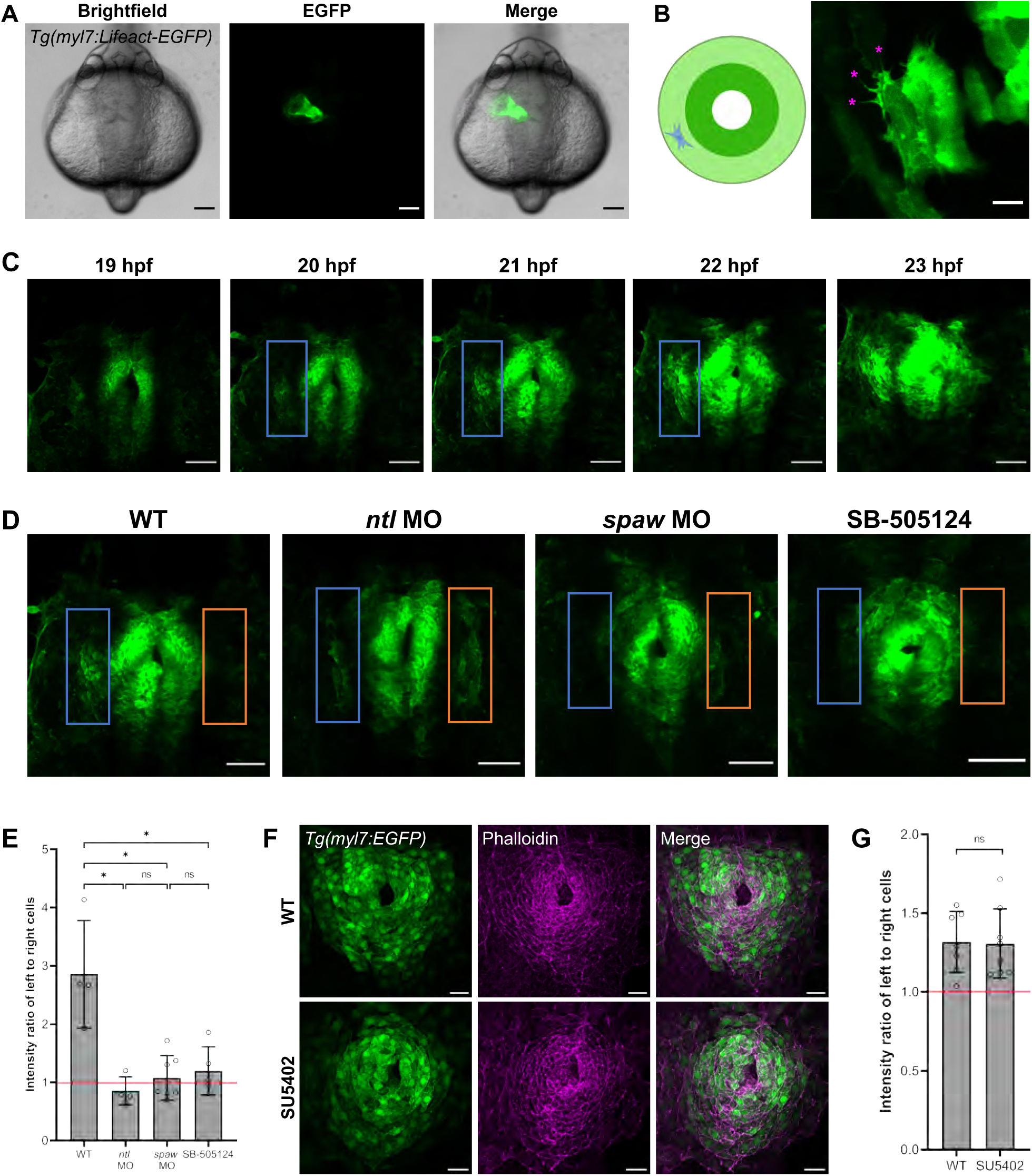
F-actin dynamics are asymmetric and Nodal-dependent in CPCs during jogging. (**A**) *Tg(myl7:Lifeact-EGFP)* embryo at 24 hpf demonstrating the fluorescence of the CPCs composing the heart tube. Scale = 100 µm. (**B**) F-actin protrusions (pink asterisks) in a singular CPC of a mosaic *Tg(myl7:Lifeact-EGFP)* embryo at 21 hpf (right image). Schematic (left image) shows the location of the CPC in the cardiac cone. Scale = 2.5 µm. (**C**) F-actin protrusive activity in left (blue box) compared to right CPCs in the cardiac cone of a WT *Tg(myl7:Lifeact-EGFP)* embryo throughout jogging from 19-23 hpf. Scale = 50 µm. (**D**) Cardiac cones of *Tg(myl7:Lifeact-EGFP)* WT, *ntl* morphant, *spaw* morphant, and SB505124-treated embryos, with left-sided (blue box) and right-sided (orange box) atrial CPCs outlined. Scale = 50 µm. (**E**) Fluorescence intensity ratio between left-sided and right-sided CPCs in the *Tg(myl7:Lifeact-EGFP)* embryos. n is WT = 4, *ntl* MO = 4, *spaw* MO = 7, SB-505124 = 5; Student’s t-test. (**F**) Cardiac cones of *Tg(myl7:EGFP)* WT and SU5402-treated embryos stained with phalloidin. Scale = 25 µm. (**G**) Fluorescence intensity ratio between left-sided and right-sided CPCs in the phalloidin-stained embryos. n is WT = 7, SU5402 = 8; Student’s t-test. (**A-D, F**) Images are dorsal views, with anterior to the top and left to the reader’s left. ns = not significant, * = p<0.01.

To determine how the actin dynamics in CPCs respond to Nodal signaling, we examined F-actin dynamics in WT embryos and compared these dynamics to those of *ntl* morphants, *spaw* morphants, and embryos treated with SB-505124, a pharmacological inhibitor of Nodal signaling (DaCosta Byfield et al., 2004; Hagos and Dougan, 2007). In WT embryos, we find a stronger Lifeact/F-actin signal in left-sided atrial cells of the cardiac cone compared to the corresponding right-sided cells, and this asymmetry persists throughout jogging (Fig. 4C, Movie 7). Left-right asymmetry in F-actin is lost when Nodal signaling is absent in *spaw* morphants or with administration of SB-505124. In fact, there was minimal F-actin signal observed in the atrial cells of these embryos (Fig. 4D, Movies 8-9). Conversely, we observe enrichment of F-actin in both left and right atrial cells in *ntl* morphants, in which Nodal is expressed bilaterally in the LPM (Fig. 4D, Movie 10). To quantify these results, we measured the ratio of fluorescence intensity values between left-sided and right-sided atrial cells (Fig. 4E). An intensity ratio of left-to-right cells greater than 1 is indicative of left-biased levels, while an intensity ratio equal to 1 indicates symmetric levels. We found the fluorescence intensity ratio in WT embryos to be left-biased, which is indicative of asymmetric F-actin levels (Fig. 4E; Table S1). However, *ntl* morphants, *spaw* morphants, and SB505124-treated embryos had ratios indicative of symmetric F-actin levels (Fig. 4E). Taken together, these findings suggest that asymmetries in Nodal correlate with asymmetries in F-actin formation in migrating atrial CPCs, suggesting a possible mechanism by which Nodal could induce left-right migration asymmetries in CPCs.

Given FGF inhibition reduces CPC migration velocity, we hypothesized this could also be due to modulation of the actin cytoskeleton. We therefore treated *Tg(myl7:EGFP)* embryos with SU5402, stained them with phalloidin, and imaged the resulting cardiac cones. We find that the left-to-right intensity ratio is left-biased in WT and SU5402 embryos (Fig. 4F-G, Table S2). These ratios are not significantly different from one another, suggesting that F-actin asymmetries are not influenced by FGF. This supports our previous data that FGF does not influence left-right cardiac laterality.

### Proper cardiac looping requires FGF induction of the second heart field

FGF signaling is known to influence various aspects of zebrafish heart morphogenesis, such as CPC differentiation and proliferation, as well as chamber size and identity establishment (de Pater et al., 2009; Felker et al., 2018; Lazic and Scott, 2011; Marques et al., 2008; Pradhan et al., 2017; Reifers et al., 1998; Zeng and Yelon, 2014). In embryos treated with SU5402 from 24-48 hpf (continuous treatment), differences in heart morphology between treated and WT embryos are apparent, particularly defects in heart looping (Fig. 5A-B). Dextral heart looping is a highly conserved process in vertebrates where the heart tube bends to place the chambers of the heart in their final, correct positions. To determine if FGF signaling influences looping at a time point beyond jogging, we treated embryos with SU5402 during specific developmental windows: during the heart tube elongation phase (24-30 hpf, prior to the onset of looping) and the heart tube bending phase (30-36 hpf, during looping) and observed the resulting effects on hearts at 48 hpf (Fig. 5A-B). Given that FGF signaling did not influence cardiac jogging laterality, we suspected that it would not influence looping laterality, as proper lateralization of the heart during jogging promotes proper looping (Grimes et al., 2020). Accordingly, embryos treated with SU5402 at either time point looped dextrally as expected (Fig. 5C).

**Figure 5.**
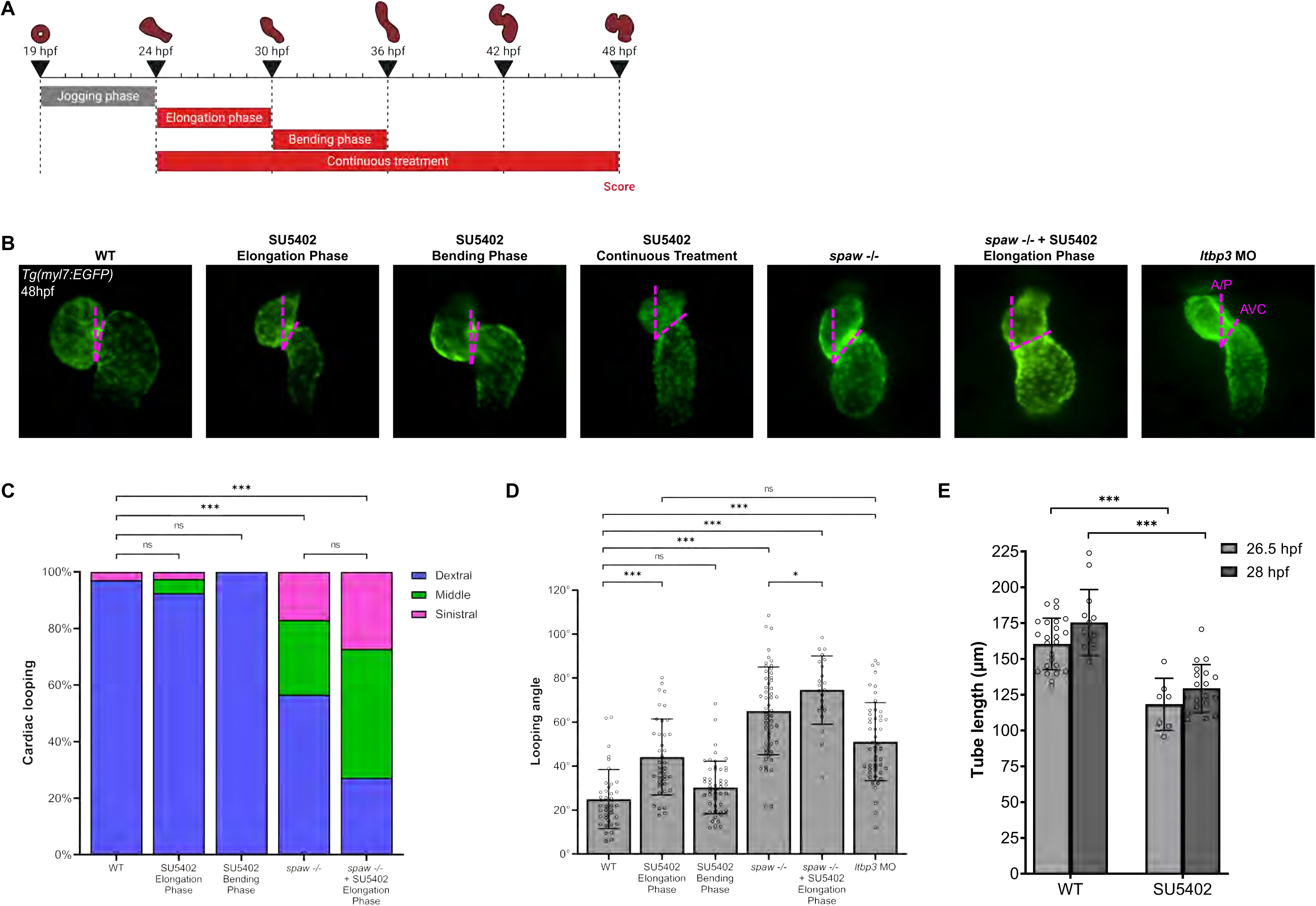
FGF signaling is necessary for SHF addition and proper cardiac looping. (**A**) Schematization depicting the SU5402 treatment windows during the “Elongation phase” from 24-30 hpf, the “Bending phase” from 30-36 hpf, and the “Continuous treatment” window from 24-48 hpf. (**B**) Representative images of WT, SU5402-treated, *spaw^-/-^,* and *ltbp3* morphant *Tg(myl7:EGFP)* embryo hearts at 48 hpf, depicting how the looping angle is measured (pink lines; AP = anterior/poster axis line and AVC = atrioventricular canal). (**C**) Quantification of looping laterality phenotypes. n is WT = 70, SU5402 elongation phase = 40, SU5402 bending phase = 50, *spaw^-/-^* = 53, *spaw^-/-^* + SU5402 elongation phase = 22; Chi-square analysis. (**D**) Quantification of looping angle. n is WT = 38, SU5402 elongation phase = 39, SU5402 bending phase = 50, *spaw^-/-^* = 53, *spaw^-/-^*+ SU5402 elongation phase = 22, *ltbp3* MO = 50; Student’s t-test. (**E**) Quantification of heart tube length in WT and SU5402-treated *Tg(nkx2.5:ZsYellow)* embryo heart tubes at 26.5 or 28 hpf. n is WT 26.5 hpf = 24, SU5402 26.5 hpf = 7, WT 28 hpf = 13, SU5402 28 hpf = 19; Student’s t-test. (**B**) Images are ventral views, with anterior to the top and left to the reader’s right. ns = not significant, * = p<0.05, *** = p<0.0001.

Quantification of successful looping can be accomplished by measuring the looping angle: the angle formed by the intersection of a line along the anterior-posterior axis and a line through the atrioventricular canal (AVC) at 48 hpf (Fig. 5B), with a properly developed heart having a looping angle of ∼25.0° (Chernyavskaya et al., 2012). In WT embryos, the heart completes looping and the atrium and ventricle are correctly positioned side-by-side at 48 hpf (Fig. 5B). Quantification of looping angle in WT *Tg(myl7:EGFP)* embryos yields a mean looping angle of 25.0° (Fig. 5D). Despite not altering looping laterality, SU5402 treatment did result in errors in looping angle. Treatment with SU5402 during the elongation phase results in significantly larger looping angles, suggesting looping is impaired when FGF signaling is inhibited prior to the start of looping (Fig. 5B, 5D). Treatment during the bending phase does not affect looping angles, indicating that inhibition of FGF signaling during the start of looping does not impair the process (Fig. 5B, 5D). Heart looping is compromised in *spaw* mutants, as expected given Nodal is known to be necessary for robust looping (Grimes et al., 2020; Noel et al., 2013) (Fig. 5B, 5D). We found that loss of FGF signaling during the elongation phase in *spaw* mutants exacerbated heart looping defects as quantified by looping angle (Fig. 5B, 5D). Our results suggest that Nodal and FGF synergize to promote heart looping.

It is known that FGF signaling is necessary for the differentiation and addition of the SHF to the arterial pole of the heart tube in zebrafish post-24 hpf (de Pater et al., 2009; Felker et al., 2018; Lazic and Scott, 2011). We hypothesized that inhibition of FGF from 24 to 30 hpf in our study may prevent development of the SHF, which may in turn influence heart looping. To test our hypothesis, we treated *Tg(nkx2.5:ZsYellow)* embryos with SU5402 during the elongation phase and measured the elongating heart tube at both 26.5 and 28 hpf. *Tg(nkx2.5:ZsYellow)* marks both the first and second heart fields, and absence of the SHF can be visualized as a shorter heart tube (Nevis et al., 2013). Measuring heart tube length revealed significantly shorter heart tubes in SU5402-treated embryos at both time points, likely due to the loss of the SHF (Fig. 5E).

To further test whether the effect on heart looping we observe is due to loss of the SHF, we inhibited SHF development by injecting embryos with a morpholino against *ltbp3* as described (Zhou et al., 2011). Similar to embryos treated with SU5402 from 24-30 hpf, *ltbp3* morphants exhibit comparable defects in looping angle (Fig. 5B, 5D). Taken together, these data suggest that inhibiting FGF signaling from 24-30 hpf affects proper cardiac looping through loss of the SHF.

## Discussion

### Nodal and FGF signals function synergistically to promote migration underlying cardiac jogging and tube extension

Our results show that both Nodal and FGF signals promote overall cell migration during heart tube formation and asymmetric placement. Losing either signal reduces cell velocities in the cone and delays the cone-to-tube transition compared to WT, while losing both signals more severely perturbs CPC migration and tube formation. During jogging, our data suggests that FGF signals act permissively to promote motility, while Nodal signals act instructively to direct CPC laterality (Fig. 6). FGF signaling is known to cooperate with Nodal signaling in coupling cell migration in various morphogenetic events, including in the *Ciona* neural tube and the zebrafish brain (Navarrete and Levine, 2016; Regan et al., 2009; Roussigne et al., 2018). Intriguingly, in the zebrafish brain, FGF and Nodal signaling function together to control left-right asymmetry by mediating collective cell migration; Nodal restricts and biases the activation of FGF to leading parapineal cells, resulting in a leftward, asymmetric migration (Regan et al., 2009; Roussigne et al., 2018). FGF is critical for parapineal cell migration, but Nodal signaling directs the laterality of this movement. This is in line with our findings that FGF is critical for overall CPC migration while Nodal directs the laterality of CPC movements.

**Figure 6.**
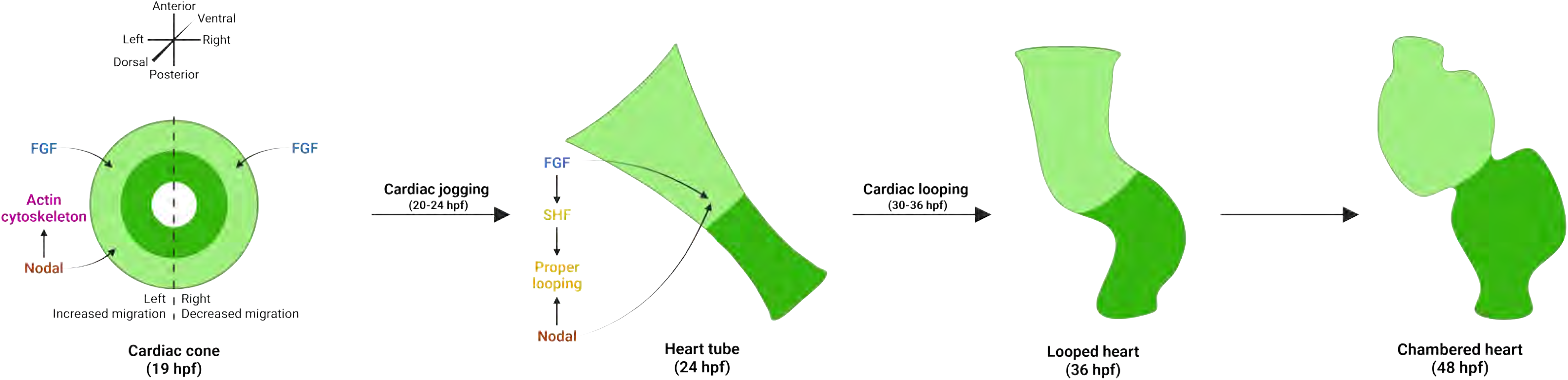
Model of the synergistic and distinct roles of Nodal and FGF signaling in asymmetric cardiac morphogenesis. In the cardiac cone, Nodal promotes a left-lateralized migration to generate cardiac asymmetry, while FGF promotes a permissive migration in all CPCs. Nodal signaling also impinges on the actin cytoskeleton to promote F-actin asymmetry. Next, in the heart tube, both Nodal and FGF signals promote proper cardiac looping. Nodal promotes looping laterality to correctly position the chambers, while FGF induces the SHF which, in turn, is likely necessary for robust looping. The looped heart then matures to form a two-chambered heart by 48 hpf.

We also find that Nodal and FGF synergize to affect the length of the heart tube at 24 hpf following jogging. Whether this defect is only the consequence of compromised motility, or whether there are other forces that restrict elongation, remains to be determined. For example, the defects we observe are highly reminiscent of *heart and mind* (*had*) and *heart and soul* (*has*) mutants, which have reduced heart tube extension (Rohr et al., 2006; Shu et al., 2003). Interestingly, *had* and *has* encode regulators of epithelial polarity, the establishment and maintenance of which is crucial for proper cardiac morphogenesis. Interrogating epithelial polarity and cell shape changes under the conditions in our study may prove informative in understanding heart tube extension. Alternatively, the decrease in heart tube length we observe in our study conditions could be due to tissue constraints on the tube as it attempts to elongate closer to the midline. Future studies are needed to determine what drives the overall anterior displacement of the heart cone/tube and the forces that drive tube extension during development.

### Nodal signals induce left-right asymmetries in F-actin that correlate with asymmetric migration

We demonstrate that Nodal signaling induces an increase in F-actin levels in migrating left atrial cells that is not observed in right atrial cells. This increase in F-actin is Nodal dependent, as it is lost in embryos injected with *spaw* morpholino or treated with Nodal inhibitor. In agreement, in embryos injected with *ntl* morpholino, where Nodal signals to both sides of the cardiac cone, both left and right atrial cells show increases in F-actin. This suggests that Nodal induces asymmetric migration by increasing actin dynamics within left atrial cells, which migrate faster than their counterparts on the right, driving clockwise rotation of the cone. Previous studies have suggested roles for Nodal in inducing cytoskeletal rearrangements in migrating cells (Noel et al., 2013; Schier, 2009; Woo et al., 2012) lending support to our hypothesis. Given heart laterality is unaltered in SU5402-treated embryos, we also hypothesized that F-actin asymmetry would not be lost in FGF-deficient embryos. We indeed found this to be the case in our phalloidin-stained SU5402-treated embryos, whose actin cytoskeleton asymmetry did not significantly differ from WT. Future studies can delve further into how loss of FGF affects CPC migration at the cellular level.

The cells in the cone are epithelial and migrate collectively during jogging. Intriguingly, using mosaic labeling with our Lifeact reporter, we observed highly dynamic, protrusive F-actin activity in these left-sided migrating cells. It is intriguing to speculate that these are “cryptic” basal extensions that are driving migration similar to what has been observed in cultured MDCK kidney cells during wound healing (Farooqui and Fenteany, 2005). Future studies can determine if these protrusions are basal-to-junctional complexes coming only from edge cells or if additional cells deeper in this epithelium are helping drive migration.

### FGF signaling is necessary for proper cardiac looping

Extending our studies with the FGF inhibitor SU5402, we find that treatment with SU5402 from 24 to 30 hpf (during heart tube elongation) affected cardiac looping and final chamber placement, while treatment from 30 to 36 hpf (during heart tube looping) did not affect looping (Fig. 6). This suggests that FGF continues to play important roles in asymmetric heart morphogenesis after jogging is complete. Cardiac looping is a highly conserved event in vertebrate heart development, but much remains to be discovered regarding its underlying mechanisms. We previously reported that jogging correctly increases the correct dextral loop in the heart, suggesting that the first asymmetric migration at 20 hpf still provides input to heart-intrinsic chirality mechanisms at later stages (Grimes et al., 2020). Our work here suggests that at least one additional input into looping *in vivo* involves FGF. We provide evidence that this input may be through SHF addition, given the similarity in phenotypes between *ltbp3* morphants and SU5402-treated embryos, both of which lack SHF structures. This is further supported by work showing heart looping defects in *isl2a* and *isl2b* mutant embryos, which produce defects in the SHF (Witzel et al., 2017). Biomechanical modeling also suggests that it is the rotation of the poles of the heart tube that drives looping, and a lack of SHF would prevent this rotation from occurring properly (Manner, 2004). Determining how the SHF contributes to proper heart looping in zebrafish will be an important area for future studies. Intriguingly, we show that Nodal and FGF both promote heart looping, as losing both pathways produces more severe defects in this process than losing either alone.

### Concluding remarks and future directions

Here, we highlight molecular mechanisms through which the spatiotemporal dynamics of signaling cues influence cardiac progenitor cell behaviors and heart morphogenesis. We further show that the differential effects exerted by interacting signals on CPCs manifest as dramatic asymmetries in heart tube morphogenesis, highlighting the fact that earlier patterning events regulate morphogenetic processes occurring later in development.

Our study suggests areas of future research on how signaling pathways synergize to promote morphogenesis. In addition to the ideas already discussed, we note that while jogging was most significantly delayed in embryos lacking both Nodal and FGF, heart tubes nevertheless ultimately formed, raising the question of what additional signals govern tube formation under these conditions. We also don’t have evidence as to whether the effect of FGF signaling on CPC migration is direct or indirect. We hypothesize it will be direct given we and others have shown that FGF components, like *fgf8*, are expressed in the early heart. Transgenic lines that modulate FGF signaling specifically in the myocardium could provide clues as to how FGF influences the CPCs during jogging and looping.

We note that the decreased velocity we observe here in *ntl* mutants conflicts with our previously published data using *ntl* morphants, in which we observed higher velocities in CPCs compared to WT (Lenhart et al., 2013). It is possible that there may be an unexpected difference in *ntl* mutants versus morphants that produces this discrepancy (Stainier et al., 2017). However, distinct microscopes, acquisition parameters, and analysis software were utilized in this study could also account for differences in velocity values, compared to our previous publication.

Ultimately, further interrogation of the roles of Nodal and FGF signaling throughout heart morphogenesis will prove informative in understanding how signaling and cellular aberrancies manifest as morphological defects. Given heart tube malformations are among the most commonly diagnosed congenital heart defects, knowing how they arise is crucial for reducing the mortality and morbidity associated with CHDs.

## Materials and methods

### Zebrafish strains

All experimental procedures in this study were conducted in accordance with the Princeton University Institutional Animal Care and Use Committee. Zebrafish (*Danio rerio*) embryos were maintained at 28°C and grown in E3 embryo medium (5 mM NaCl, 0.17 mM KCl, 0.33 mM CaCl_2_, 0.33 mM MgSO_4_). We used wild-type strains WIK and PWT. We used the zebrafish mutants *ntl^b160^* and *spaw^sa177^* (Grimes et al., 2020; Halpern et al., 1993). We used the transgenic strains *Tg(myl7:EGFP)twu34* (Huang et al., 2003)*, TgBAC(-36nkx2.5:ZsYellow)fb7* (Zhou et al., 2011), and *Tg(hsp70l:dnfgfr1a-EGFP)pd1;* (Lee et al., 2005).

### Generation of transgenic line

We generated the *Tg(myl7:Lifeact-EGFP)pr26* transgenic strain using the Tol2 transposase system (Kawakami, 2007). Briefly, 1-cell stage embryos were injected with Tol2 mRNA and *pTol2005b-myl7:Lifeact-EGFP* plasmid. Only correctly developing embryos with EGFP-positive hearts were raised. Injected F_0_ fish were outcrossed to generate stable transgenic lines.

### Drug treatments

A 10 mM stock of the FGFR inhibitor SU5402 (Sigma-Aldrich) in DMSO was diluted to a working concentration of 6 μM or 10 μM in a 1% DMSO/1X E3 solution. 6 μM was used for inhibition during bending and elongation phases and 10 μM was used for inhibition during jogging phase. Embryos were dechorionated and incubated at 28°C in the dark for varying durations of time: 19-24 hpf, 24-30 hpf, 30-36 hpf, and 24-48 hpf. In our experiments, expression of the FGF signaling targets *pea3* and *erm* is lost within 30 minutes of treatment initiation. Therefore, addition of SU5402 occurred 30 minutes before each inhibition phase. Upon completion of SU5402 incubation, embryos were either rinsed with E3 and raised to the desired stage, underwent live imaging, or fixed in 4% paraformaldehyde (PFA; Electron Microscopy Sciences) for RNA *in situ* analysis. Due to the light-sensitivity of SU5402, treated embryos were incubated in the dark. For SB-505124 (Sigma-Aldrich) treatments, a 10 mM stock in DMSO was diluted to a working concentration of 40 μM and administered to embryos at the tailbud stage; embryos then underwent live imaging during jogging as described (Lenhart et al., 2013). Note that DMSO-treated vehicles did not differ from WT embryos in the processes studied.

### Heat shock treatment

*Tg(hsp70l:dnfgfr1-EGFP)pd1* embryos underwent heat shock at 38°C for 30 minutes every 2 hours from 17 hpf until fixation to ensure FGF inhibition during jogging phase. The dominant negative FGF construct has been found to activate approximately 2 hours after initial heat shock (Lee et al., 2014; Pouget et al., 2014; Vemaraju et al., 2012). Thus, starting heat shock at 17 hpf allowed for FGF inhibition in the same time window as in our experiments with SU5402. EGFP expression served as confirmation of successful heat shock. Hearts at 26.5 hpf appeared comparable to WT in shape and size, indicating CPC differentiation was not affected by treatment during this timeframe.

### RNA and morpholino injections

psMEK mRNA injections were performed as previously described (Patel et al., 2019). Briefly, 50 pg of psMEK^E203K^ mRNA was injected into the yolk at the 1-cell stage. Injected embryos were immediately placed in a dark box until 18.5 hpf. The lid of the box was then removed and replaced with an LED board emitting 505 nm light from 18.5 to 24 hpf to activate MEK signaling during jogging phase. Morpholino injections were performed as previously described using established morpholinos for *spaw*, *ntl*, and *ltbp3* (Lenhart et al., 2013; Long et al., 2003; Nasevicius and Ekker, 2000; Zhou et al., 2011). Note that control embryos injected with phenol red and nuclease-free water for both psMEK and morpholino injections did not differ from WT embryos in the processes studied.

### RNA *in situ* hybridization

Chromogenic whole-mount RNA *in situ* hybridization was performed using digoxygenin-labeled RNA probes for *myl7* and *lefty2* (Berdougo et al., 2003; Bisgrove et al., 1999; Thisse and Thisse, 2014; Yelon et al., 1999). Images were acquired using the Leica DMRA microscope. Fluorescent whole-mount *in situ* hybridization was performed using the Molecular Instruments HCR kit as described (Choi et al., 2018) for probes against *myl7* and *fgf8*. Images were acquired using the Nikon A1 confocal microscope.

### Confocal imaging and analysis

To analyze cell migration during cardiac jogging, dechorionated embryos were mounted in 0.8% low-melt agarose as previously described (Lenhart et al., 2013). The dish was covered with a 0.5% DMSO/1X E3 solution containing 0.13 mM tricaine (MS-222; Sigma-Aldrich). For SU5402 treatments, a 5 μM dilution in 0.8% agarose was used for mounting embryos, which were incubated in inhibitor at least 30 minutes before mounting and covered with 10 μM inhibitor in 0.5% DMSO/1X E3 containing 0.13 mM tricaine.

For analysis of cardiac cell movements, embryos were imaged from 18-24 hpf using a Leica SP5 confocal microscope. The Leica Mark & Find feature was used to image up to 4 embryos per imaging session for higher throughput. A heated stage set to 28.5°C was used, which permitted imaging for several hours without perturbing development. We acquired confocal z-stacks at 4- or 5- minute intervals with a spacing of 2 μm.

To track individual cells during jogging, we compressed the z-stacks into a single dimension and then utilized the ImageJ program TrackMate to track and quantify the metrics of each cell during its migration. To ensure consistent quantification of velocity over time between different genotypes, we tracked CPCs over the same developmental window: we began tracking cells upon anterior fusion of CPCs at the midline and ended tracking 3 hours thereafter. Note that only outer, atrial cells were chosen for tracking as they undergo the most migration during jogging; the bright ventricular cells around the lumen are excluded from analysis (Lenhart et al., 2013). Average velocity and angle of displacement for each cell were calculated using raw position, displacement, and time measurements generated by TrackMate. Resulting cell velocities, angles, and displacements were input into Prism or RStudio for analysis. Note that distinct microscopes, acquisition parameters, and analysis software were utilized and may account for differences in velocity values compared to our previous publications.

For analysis of F-actin in *Tg(myl7:Lifeact-EGFP)pr26* embryos, embryos were mounted and handled as described above. Embryos were imaged from 19-23 hpf using a Prairie Ultima two-photon microscope. Z-stacks were acquired at 15*-*minute intervals with a spacing of 4 μm. Fluorescence intensity analysis was performed using ImageJ.

For analysis of F-actin via phalloidin in *Tg(myl7:EGFP)* embryos, embryos were dechorionated and fixed in 4% PFA with 4% sucrose and fixed overnight at 4°C. Embryos were then washed three times with PBS, washed three times with PBS-Triton, and then stained overnight with 1:50 phalloidin rhodamine (Cytoskeleton Inc. #PHDR1) at 4°C. Embryos were then washed four times with PBS-Triton and transferred to 30% glycerol and then 50% glycerol in PBS. Embryos were then deyolked, mounted, and imaged using a Nikon A1R-Si confocal microscope. The 40X objective was used, and Z-stacks had a spacing of 1 μm. Fluorescence intensity analysis was performed using ImageJ.

For observing cardiac looping, embryos were mounted and handled as described above. Images were acquired using the Leica M205 FA fluorescent stereoscope.

### Heart measurements and scoring

For observing heart tube length in fixed embryos processed via RNA *in situ* hybridization, embryos were observed using the Leica S6E dissecting microscope, and heart tube length was measured through the eyepiece using an ocular micrometer and a stage micrometer. For observing heart tube length and cardiac looping in live embryos, embryos were mounted and handled as described above. Images were acquired using the Leica M205 FA fluorescent stereoscope. Analysis was performed using ImageJ ruler and angle tools. The looping angle is defined as the angle between the plane of the anterior-posterior axis and the AVC (Chernyavskaya et al., 2012).

For scoring of heart laterality, embryos were staged at 26.5 hpf and scored as previously described (Grimes et al., 2020; Lenhart et al., 2013). Briefly, the heart tubes of embryos were viewed dorsally using a stereomicroscope. A heart tube that remained along the midline was scored as midline; a tube that deviated from the midline was scored as having jogged left or right.

For measuring fluorescent intensity in the cardiac cone via ImageJ, the atrial cells of the left or right sides were outlined, and the measure function was used to quantify the mean gray value. The left value was divided by the right value to yield the left:right fluorescence intensity ratio.

### Statistical analysis

All bar graphs were generated using GraphPad Prism 9.4.0. Statistical tests of significance (Student’s t-test, Chi-square analysis) were calculated using GraphPad Prism analysis software. Note at least N = 2 biological and technical replicates were performed for each experiment.

## Acknowledgments

We would like to thank Drs. Stefano Di Talia, Fang Lin, C. Ben Lovely, Tatjana Piotrowski and Kenneth Poss for providing us with zebrafish transgenic strains to conduct this work. We thank Dr. Gary Laevsky and the Confocal Imaging Facility, a Nikon Center of Excellence, in the Department of Molecular Biology for their assistance with imaging. We also thank Phillip Johnson and his team for their assistance with fish husbandry.

## Data and resource availability

All relevant data and resources can be found within the article and its supplementary information. The data that support the findings of this study are available from the corresponding author RDB, upon reasonable request.

## Diversity and inclusion statement

One or more of the authors of this paper self-identifies as an underrepresented ethnic minority in science. One or more of the authors of this paper self-identifies as living with a disability. One or more authors of this paper received support from a program designed to increase minority representation in science. Five of the six authors on this paper are women, who are often underrepresented in science.

