## Supplementary Information for "Synergistic and independent roles for Nodal and FGF in zebrafish CPC migration and asymmetric heart morphogenesis"

**Movie 1.** Live imaging of CPCs jogging in a WT transgenic *Tg(myl7:EGFP)* embryo from the onset of cardiac cone formation until three hours after.

**Movie 2.** Live imaging of CPCs jogging in a *ntl* mutant *Tg(myl7:EGFP)* embryo from the onset of cardiac cone formation until three hours after.

**Movie 3.** Live imaging of CPCs jogging in a *spaw* mutant *Tg(myl7:EGFP)* embryo from the onset of cardiac cone formation until three hours after.

**Movie 4.** Live imaging of CPCs jogging in a SU5402-treated *Tg(myl7:EGFP)* embryo from the onset of cardiac cone formation until three hours after.

**Movie 5.** Live imaging of CPCs jogging in a *spaw* mutant and SU5402-treated *Tg(myl7:EGFP)* embryo from the onset of cardiac cone formation until three hours after.

**Movie 6.** Live imaging of F-actin in a singular cardiac progenitor cell (CPC) from a WT mosaic *Tg(myl7:Lifeact-EGFP)* embryo at 21 hours post fertilization (hpf). Movie is a total of 5 minutes.

**Movie 7.** Live imaging of F-actin in a WT transgenic *Tg(myl7:Lifeact-EGFP)* embryo from 19-23 hpf, following the cardiac cone as it undergoes jogging.

**Movie 8.** Live imaging of F-actin in a *spaw* morphant *Tg(myl7:Lifeact-EGFP)* embryo from 19-23 hpf, following the cardiac cone as it undergoes jogging.

**Movie 9.** Live imaging of F-actin in a SB-505124-treated *Tg(myl7:Lifeact-EGFP)* embryo from 19-23 hpf, following the cardiac cone as it undergoes jogging.

**Movie 10.** Live imaging of F-actin in a *ntl* morphant *Tg(myl7:Lifeact-EGFP)* embryo from 19-23 hpf, following the cardiac cone as it undergoes jogging.

**Table S1.** Fluorescence intensity values of *Tg(myl7:Lifeact-EGFP)* embryos.

| Embryo | Condition | Side | Mean Fluorescence Value | Ratio (L/R) |
|---|---|---|---|---|
| 1 | WT | L | 1349.94 | 2.69 |
| 1 | WT | R | 501.77 | |
| 2 | WT | L | 2322.77 | 1.94 |
| 2 | WT | R | 1199.43 | |
| 3 | WT | L | 621.93 | 2.67 |
| 3 | WT | R | 233.15 | |
| 4 | WT | L | 2220.61 | 4.14 |
| 4 | WT | R | 536.99 | |
| 5 | ntl MO | L | 547.29 | 0.77 |
| 5 | ntl MO | R | 715.64 | |
| 6 | ntl MO | L | 692.61 | 0.79 |
| 6 | ntl MO | R | 879.54 | |
| 7 | ntl MO | L | 645.13 | 0.67 |
| 7 | ntl MO | R | 968.04 | |
| 8 | ntl MO | L | 1123.93 | 1.20 |
| 8 | ntl MO | R | 934.84 | |
| 9 | SB-505124 | L | 811.40 | 0.96 |
| 9 | SB-505124 | R | 845.70 | |
| 10 | SB-505124 | L | 642.86 | 0.81 |
| 10 | SB-505124 | R | 797.09 | |
| 11 | SB-505124 | L | 1382.32 | 1.32 |
| 11 | SB-505124 | R | 1045.72 | |
| 12 | SB-505124 | L | 958.35 | 1.04 |
| 12 | SB-505124 | R | 918.12 | |
| 13 | SB-505124 | L | 509.50 | 1.86 |
| 13 | SB-505124 | R | 274.27 | |
| 14 | spaw MO | L | 766.05 | 0.80 |
| 14 | spaw MO | R | 953.62 | |
| 15 | spaw MO | L | 3281.47 | 1.71 |
| 15 | spaw MO | R | 1916.12 | |
| 16 | spaw MO | L | 466.68 | 0.83 |
| 16 | spaw MO | R | 564.29 | |
| 17 | spaw MO | L | 1062.37 | 1.37 |
| 17 | spaw MO | R | 774.68 | |
| 18 | spaw MO | L | 645.79 | 0.70 |
| 18 | spaw MO | R | 928.29 | |
| 19 | spaw MO | L | 417.44 | 1.30 |
| 19 | spaw MO | R | 320.33 | |
| 20 | spaw MO | L | 401.35 | 0.84 |
| 20 | spaw MO | R | 476.79 | |

**Table S2.** Fluorescence intensity values of phalloidin in *Tg(myl7:EGFP)* embryos.

| Embryo | Condition | Side | Mean Fluorescence Value | Ratio (L/R) |
|---|---|---|---|---|
| 1 | WT | L | 270.82 | 1.55 |
| 1 | WT | R | 174.59 | |
| 2 | WT | L | 349.40 | 1.49 |
| 2 | WT | R | 234.08 | |
| 3 | WT | L | 329.57 | 1.23 |
| 3 | WT | R | 267.16 | |
| 4 | WT | L | 222.92 | 1.04 |
| 4 | WT | R | 214.63 | |
| 5 | WT | L | 262.56 | 1.29 |
| 5 | WT | R | 203.26 | |
| 6 | WT | L | 278.47 | 1.14 |
| 6 | WT | R | 244.65 | |
| 7 | WT | L | 244.45 | 1.47 |
| 7 | WT | R | 166.11 | |
| 8 | SU5402 | L | 136.55 | 1.12 |
| 8 | SU5402 | R | 121.94 | |
| 9 | SU5402 | L | 196.49 | 1.71 |
| 9 | SU5402 | R | 114.59 | |
| 10 | SU5402 | L | 393.22 | 1.13 |
| 10 | SU5402 | R | 349.39 | |
| 11 | SU5402 | L | 383.40 | 1.31 |
| 11 | SU5402 | R | 293.20 | |
| 12 | SU5402 | L | 338.85 | 1.34 |
| 12 | SU5402 | R | 251.94 | |
| 13 | SU5402 | L | 336.59 | 1.11 |
| 13 | SU5402 | R | 303.46 | |
| 14 | SU5402 | L | 390.03 | 1.53 |
| 14 | SU5402 | R | 254.55 | |
| 15 | SU5402 | L | 338.41 | 1.20 |
| 15 | SU5402 | R | 283.11 | |
